# Accounting for 16S rRNA copy number prediction uncertainty and its implications in bacterial diversity analyses

**DOI:** 10.1101/2021.08.31.458422

**Authors:** Yingnan Gao, Martin Wu

## Abstract

16S rRNA gene copy number (16S GCN) varies among bacterial species and this variation introduces potential biases to microbial diversity analyses using 16S rRNA read counts. To correct the biases, methods have been developed to predict 16S GCN. A recent study suggests that the prediction uncertainty can be so great that copy number correction is not justified in practice. Here we develop RasperGade16S, a novel method and software to better model and capture the inherent uncertainty in 16S GCN prediction. RasperGade16S implements a maximum likelihood framework of pulsed evolution model and explicitly accounts for intraspecific GCN variation and heterogeneous GCN evolution rates among species. Using cross validation, we show that our method provides robust confidence estimates for the GCN predictions and outperforms other methods in both precision and recall. We have predicted GCN for 592605 OTUs in the SILVA database and tested 113842 bacterial communities that represent an exhaustive and diverse list of engineered and natural environments. We found that the prediction uncertainty is small enough for 99% of the communities that 16S GCN correction should improve their compositional and functional profiles estimated using 16S rRNA reads. On the other hand, we found that GCN variation has limited impacts on beta-diversity analyses such as PCoA, PERMANOVA and random forest test.

## Introduction

The 16S ribosomal RNA (16S rRNA) gene is the gold standard for bacterial and archaeal diversity study and has been commonly used to estimate the composition of bacterial and archaeal communities through amplicon sequencing. Sequence reads are usually matched to reference databases like SILVA [1] and GreenGenes [2] to determine the presence of taxa and their relative cell abundances. However, the 16S rRNA gene copy number (16S GCN) can vary from 1 to more than 15 [3, 4] and this large copy number variation introduces bias in the relative cell abundance estimated using the gene read counts (thereafter referred to as gene abundance) [5], and consequently it can skew the community profiles, diversity measures and lead to qualitatively incorrect interpretations [5–8]. As a result, it has been argued that 16S GCN variations should be taken into account in 16S rRNA gene-based analyses [5].

The majority of bacteria species have not been cultured or sequenced and their 16S GCNs are unknown. Studies have shown that 16S GCN exhibits a strong phylogenetic signal [5, 7], and therefore 16S GCN can be inferred from closely related reference bacteria. Based on this principle, software has been developed to predict the 16S GCN [5, 7, 9, 10] in a process often referred to as hidden state prediction [11]. However, a recent study correctly points out that the accuracy of 16S GCN prediction deteriorates as the minimum phylogenetic distance between the query sequence and the reference sequences increases, and the prediction of 16S GCN is still an open question [12].

The increasing error of 16S GCN prediction with increasing phylogenetic distance roots from the stochastic nature of trait evolution, which leads to inherent uncertainty in the predicted trait values. One way of reducing the inherent uncertainty is to improve taxon sampling in the reference phylogeny to reduce the query’s phylogenetic distance to the reference [13]. Another way of addressing the inherent uncertainty is to model the uncertainty directly and have a confidence estimate. By doing so, we will be able to determine how confident we should be about a GCN prediction and make meaningful interpretations. Unfortunately, few 16S GCN prediction tools provide a confidence estimation for the predicted 16S GCN, and uncertainty is mostly ignored when interpreting the results of downstream analyses [5, 7, 10]. For example, PICRUST2 predicts functional profiles of bacterial and archaeal communities from 16S rRNA sequence data. It predicts 16S GCN for each operational taxonomic unit (OTU) in the community and uses the predicted values (point estimates) to estimate “corrected” relative cell abundances and metagenomes, without accounting for the uncertainty of the predictions. As a result, the impact of uncertainty in 16S GCN prediction on bacterial diversity analyses remains unknown and needs to be investigated.

Several points need to be considered to properly model the prediction uncertainty. First, because the uncertainty roots from the stochastic nature of trait evolution, we need to develop a good model for 16S GCN evolution. Previously the evolution of the 16S GCN trait has been modeled as gradual evolution using the Brownian motion (BM) model [5, 7, 10]. However, alternative models exist and need to be considered [14–16]. For example, pulsed evolution (PE) model postulates that traits evolve by jumps, followed by periods of stasis [14, 17]. It has been shown that pulsed evolution is prevalent in microbial genome trait evolution [18]. 16S GCN of *Bacillus subtilis* can jump from 1 to 6 in a matter of days by gene amplification [19]. On the other hand, it is well known that the 16S GCN of some bacterial clades such as the Rickettsiales order, a diverse group of obligate intracellular bacteria, has only one copy of 16S rRNA in their genomes, demonstrating stasis [20, 21]. To develop a proper model for 16S GCN evolution, the tempo and mode of evolution need to be examined.

Secondly, 16S GCN can vary within the same species [22–25], which introduces uncertainty to GCN prediction that needs to be accounted for. It has been shown that modeling the intraspecific variation is essential for the analysis of comparative trait data and failing to account for this variation can result in model misspecification [14]. Because conspecific strains are usually separated by zero branch length in the phylogeny of the 16S rRNA gene, the intraspecific variation can be modelled as time-independent variation, which can also account for measurement errors [26].

Thirdly, there is notable rate heterogeneity in 16S GCN evolution. For example, the obligately intracellular bacteria and free-living bacteria with streamlined genomes (e.g., *Rickettsia* and *Pelagibacter*) have elevated molecular evolutionary rates [27, 28] and therefore relatively long branches in the 16S rRNA gene phylogeny [29]. Nevertheless, they have only one copy of 16S rRNA in their genomes and the GCNs rarely change [21]. It is expected that the 16S GCN prediction for this group of bacteria should be accurate despite their large phylogenetic distances to the reference genomes. Such examples suggest that the rate heterogeneity of 16S GCN evolution should be systematically evaluated and modelled properly. However, no previous methods have evaluated and modeled such evolution rate heterogeneity, leading to potential model misspecification in 16S GCN predictions.

Here, we develop a novel tool *RasperGade16S* that employs a heterogeneous pulsed evolution model for 16S rRNA GCN prediction. Through simulation and cross-validation, we show that *RasperGade16S* outperforms other methods in terms of providing significantly improved confidence estimates. We demonstrate that correcting 16S rRNA GCN improves the relative cell abundance estimates of the bacterial communities and is expected to be beneficial for more than 99% of 113842 environmental samples we analyzed. However, our findings suggest that GCN correction may not be necessary for beta-diversity analyses, as it has limited impact on the results.

## Methods

### Compiling 16S GCN data and inferring 16S rRNA reference phylogeny

We downloaded annotated RNA gene sequences from 21245 complete bacterial genomes in the NCBI RefSeq database (Release 205) on April 9, 2021. For each genome, we counted the number of annotated 16S rRNA genes. Genomes with questionable 16S GCNs were removed and one representative 16S rRNA sequence from each remaining genome was selected. A 16S rRNA phylogeny (referred to as reference phylogeny hereafter) was inferred from the representative sequences of 6408 genomes. See Supplementary Methods for details.

### Evaluating time-independent variation in 16S GCN

To evaluate the extent of 16S GCN time-independent or intraspecific variation, we compared GCN between 5437 pairs of genomes with identical 16S rRNA gene alignments. To formally test whether accounting for time-independent variation is necessary, we modeled time-independent variation as a normal white noise, and fitted the Brownian motion (BM) model to the evolution of 16S GCN in the 6408 reference genomes, with and without time-independent variation. We then calculated the likelihood and chose the best model using the Akaike Information Criterion (AIC).

### Evaluating the rate heterogeneity of 16S GCN evolution

We calculated the local average rate of evolution for each genus that contains at least 10 genomes in the reference phylogeny and examined the distribution of the average rates among genera. The average rate of a genus is calculated as the variance of phylogenetically independent contrasts (PICs) [30] of GCN within the genus.

### Predicting 16S GCN

We developed a heterogeneous pulsed evolution model to model 16S GCN evolution (see Supplementary Methods for details) and a likelihood based R package *RasperGade16S* to predict 16S GCN. *RasperGade16S* first assigns the query sequence to either the regularly-evolving or the slowly-evolving group based on where it is inserted in the reference phylogeny. For a query sequence inserted into the slowly-evolving group, its insertion branch length is scaled by the ratio *r_slow_/r_regular_*, where r is the rate of evolution in each group. For a query sequence inserted into the regularly-evolving group, a small branch length is added to the insertion branch to represent the estimated time-independent variation. *RasperGade16S* then predicts the GCN of the query using the rescaled reference phylogeny. Because 16S GCN is an integer trait, the continuous prediction from hidden state prediction is rounded and a confidence (probability) that the prediction is equal to the truth is estimated by integrating the predicted uncertainty distribution. We marked the 16S GCN prediction with a confidence smaller than 95% as unreliable, and otherwise as reliable. As a comparison, we also predicted GCN using PICRUST2, which employs multiple hidden state prediction methods in the R package *castor* [31] for 16S GCN predictions. We selected three methods by which confidence can be estimated: the phylogenetically independent contrast (pic) method, the maximum parsimony (mp) method, and the empirical probability (emp) method. Otherwise, we run PICRUST2 using default options and the unscaled reference phylogeny.

We did not test the tools CopyRighter [7] and PAPRICA [9] in this study because 1) neither provides the option of using a user-supplied reference data, and 2) neither provides uncertainty estimates (i.e., confidence intervals) of its predictions, which is the primary focus of this study.

### Adjust NSTD and NSTI with rate heterogeneity

The adjusted nearest-sequenced-taxon-distances (NSTDs) [12] is calculated using the rescaled reference tree. The adjusted nearest-sequenced-taxon-index (NSTI) [10] is calculated as the weighted average of adjusted NSTDs of the community members.

### Validating the quality of predicted 16S GCN and its confidence estimate

We used cross-validations to evaluate the quality of 16S GCN prediction and its confidence estimate, and how they vary with NSTD. We randomly selected 2% of the tips in the reference phylogeny as the test set and filtered the remaining reference set by removing tips with a NSTD to any test sequence smaller than a threshold. We then predicted the 16S GCN for each tip in the test set using the filtered reference set. We conducted cross-validation within 9 bins delineated by 10 NSTD thresholds: 0, 0.002, 0.005, 0.010, 0.022, 0.046, 0.100, 0.215, 0.464 and 1.000 substitutions/site, and for each bin we repeated the cross-validation 50 times with non-overlapping test sets. We evaluated the quality of the 16S GCN prediction by the coefficient of determination (R^2^), the fraction of variance in the true copy numbers explained by the prediction. We evaluated the quality of confidence estimate by precision and recall. Precision is defined as the proportion of accurately predicted 16S GCN in predictions considered as reliable (with ≥ 95% confidence), and recall is defined as the proportion of reliable predictions in the accurately predicted 16S GCNs. We averaged the R^2^, precision and recall for the 50 cross-validations in each bin.

### Evaluating the effect of 16S GCN correction on relative cell abundance estimation

We simulated bacterial communities with 16S GCN variation (SC1 dataset, see Supplementary Methods). To estimate the confidence interval (CI) of the corrected relative cell abundance of each OTU in a community, we randomly drew 1000 sets of 16S GCNs from their predicted uncertainty distribution. For each set of 16S GCNs, we divided the gene read count of OTUs by their corresponding 16S GCNs to get the corrected cell counts. The median of the corrected cell count for each OTU in the 1000 sets is used as the point estimate of the corrected cell count, and the OTU’s relative cell abundance is calculated by normalizing the corrected cell count with the sum of corrected cell counts of all OTUs in the community. The 95% CI for each OTU’s relative cell abundance is determined using the 2.5% and 97.5% quantiles of the 1000 sets of corrected relative cell abundances. The support value for the most abundant OTU is calculated as the empirical probability that the OTU has the highest cell abundance in the 1000 sets of corrected cell abundances. We calculated the coverage probability of the CI as the empirical frequency that the relative gene abundance or true relative cell abundance is covered by the estimated CI. We evaluated the effect of 16S GCN correction on relative cell abundance estimation at different NSTD thresholds.

### Evaluating the effect of 16S GCN correction on beta-diversity analyses

We used the Bray-Curtis dissimilarity and Aitchison distance for beta-diversity analysis that requires a dissimilarity or distance matrix and evaluated the effect of 16S GCN correction on the simulated bacterial communities (SC2 dataset, see Supplementary Methods). To correct for 16S GCN variation in beta-diversity analyses, we divided the gene abundance of each OTU by its predicted 16S GCN and calculated the corrected relative cell abundance table and the corresponding dissimilarity/distance matrix. We used the corrected cell abundance table to generate the principal coordinates analysis (PCoA) plot and to conduct the permutational multivariate analysis of variance (PERMANOVA) and the random forest test with the R package *vegan* and *randomForest*, respectively.

### Examining the adjusted NSTI of empirical bacterial communities

To check the predictability of 16S GCN in empirical data, we examined bacterial communities surveyed by 16S rRNA amplicon sequencing in the MGnify resource platform [32] that were processed with the latest two pipelines (4.1 and 5.0). The MGnify resource platform uses the SILVA database release 132 [1] for OTU-picking in their latest pipelines, and therefore we predicted GCNs for SILVA OTUs (Supplementary Methods). We filtered the surveyed communities from the MGnify platform so that only communities with greater than 80% of their gene reads mapped to the SILVA reference at a similarity of 97% or greater were included. This filtering yielded 113842 bacterial communities representing a broad range of environment types. We calculated the adjusted NSTI for each community and examined the adjusted NSTI distribution in various environmental types.

## Results

### Time-independent variation is present in 16S GCN evolution

To evaluate the extent of time-independent or intraspecific variation in 16S GCN, we examined 5437 pairs of genomes with identical 16S rRNA gene alignments. The 16S GCN differs in 607 (11%) of them, suggesting the presence of significant time-independent variation. For the 6408-genomes in the reference phylogeny, we found that incorporating time-independent variation with the BM model greatly improves the model fit (Table 1), indicating the necessity to take time-independent variation into account in 16S GCN prediction. In addition, we observed that the rate of evolution in the fitted BM model is inflated by 1670 folds when time-independent variation is not included in the model, which will lead to overestimation of uncertainty in BM model-based 16S GCN prediction.

**Table 1.** The AICs of Brownian motion model and pulsed evolution model.

| Model | BM | BM (with time-independent variation) | PE (with time-independent variation) |
| --- | --- | --- | --- |
| Homogenous model | 34338 | 18028 | -7925 |
| Heterogeneous model | NA | NA | -15395 |

### Pulsed evolution model explains the 16S GCN evolution better than the Brownian motion model

When predicting traits using phylogenetic methods, the BM model is commonly assumed to be the model of evolution. We have shown that PE model is a better model for explaining the evolution of bacterial genome size [33], prompting us to test whether pulsed evolution can be applied to explain 16S GCN evolution as well. Using the R package *RasperGade* that implements the maximum likelihood framework of pulsed evolution [14], we fitted the PE model with time-independent variation to the same dataset. Table 1 shows that the PE model provides a significantly better fit than the BM model, indicating that 16S GCN prediction should assume the PE model instead of the BM model. Fitted model parameters are not sensitive to the HMM profiles used for aligning the 16S rRNA sequences (Table S2).

### Substantial rate heterogeneity exists in 16S GCN evolution

To systematically examine the rate heterogeneity of 16S GCN evolution in the reference genomes, we first used the variance of PICs as an approximate estimate of the local evolution rate of 16S GCN. We found that the rate of evolution varies greatly among genera (Figure S1), but can be roughly divided into two groups with high and low rates of evolution. Therefore, we developed a heterogeneous pulsed evolution model where all jumps are the same size but the frequency of jumps varies between two groups to accommodate the heterogeneity among different bacterial lineages. Using a likelihood framework and AIC, we classified 3049 and 3358 nodes and their descending branches into slowly-evolving and regularly-evolving groups respectively (Figure S2). The frequency of jumps in the regularly-evolving group is 145 folds of the frequency in the slowly-evolving group (Table S3). The heterogeneous PE model provides the best fit among all models tested (Table 1), indicating that a heterogeneous PE model should be assumed in predicting 16S GCN.

Apart from the rate of pulsed evolution, we also observed heterogeneity in time-independent variation: for the slowly-evolving group, the fitted model parameters indicate no time-independent variation, while for the regularly-evolving group, the magnitude of time-independent variation is approximately 40% of a jump in pulsed evolution (Table S3). The presence of time-independent variation caps the confidence of prediction in the regularly-evolving group at 85%, which can only be achieved when the query has identical 16S rRNA gene alignment to one of the reference genomes.

### RasperGade16S improves confidence estimate for 16S GCN prediction in empirical data

Using 16S GCN from the 6408 complete genomes in the reference phylogeny for cross-validation, we compared the performance of various methods in accuracy and confidence estimates. The pic and mp methods produce very large and zero uncertainty respectively (Figure 1A), leading to both poor precision and recall rates (Figure 1C and 1D). The emp method performs the worst in terms of accuracy. The PE method produces the best overall precision (Figure 1D), achieving an average precision rate of 0.96, one of the best accuracies (Figure 1B), and the best confidence estimate for 16S GCN prediction (Figure 1C) over the full spectrum of NSTD, and should be preferred when predicting 16S GCN.

**Figure 1.**
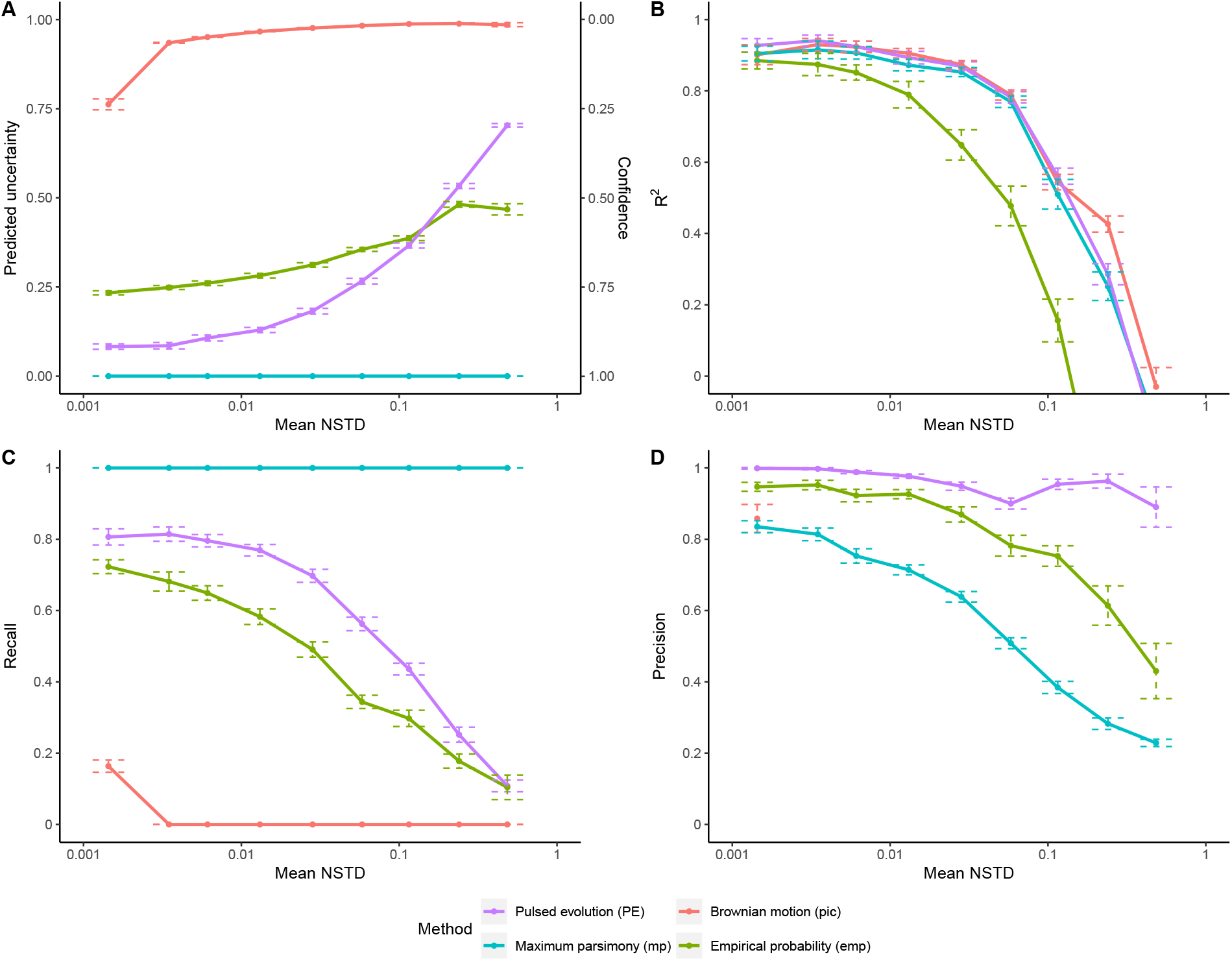
The performance of prediction on empirical 16S GCN. Using cross-validation of empirical data, the mean estimated uncertainty and confidence of predictions (A), the mean coefficient of determination R^2^ of the predictions (B), and the recall (C) and precision (D) of classification of predictions by their associated confidence estimate, plotted against the mean NSTD. The error bars represent the 95% CI of the mean. The empirical 16S GCN analyzed here are from the 6408 complete genomes in the reference phylogeny.

### Copy number correction improves relative cell abundance estimation

From theoretical calculations, in general, community members with lower relative cell abundances suffer from greater impacts by 16S GCN variation (Figure 2A). If a species has a higher GCN compared to the average GCN of the community, its relative abundance will be overestimated. Otherwise, its presence will be underestimated (Figure 2A). In simulated dataset (SC1), we found that 16S GCN variation has a large detrimental effect on the estimated relative cell abundance (Figure 2B). On average, the relative cell abundance estimated using the gene abundance increased or decreased by 1.8-fold compared to the true relative cell abundance, and the empirical probability of correctly identifying the most abundant OTU based on the gene abundance is only around 13% (Figure 2C). Correcting for 16S GCN improves the estimated relative cell abundance (Figure 2B). As expected, the improvement is greatest when the adjusted NSTI is small (i.e., when there are closely related reference genomes), and it gradually diminishes when the adjusted NSTI increases. At the smallest adjusted NSTI, the average fold change of the estimated relative cell abundance decreases to 1.1-fold after 16S GCN correction and the empirical probability of correctly identifying the most abundant OTU increases to around 65% (Figure 2C).

**Figure 2.**
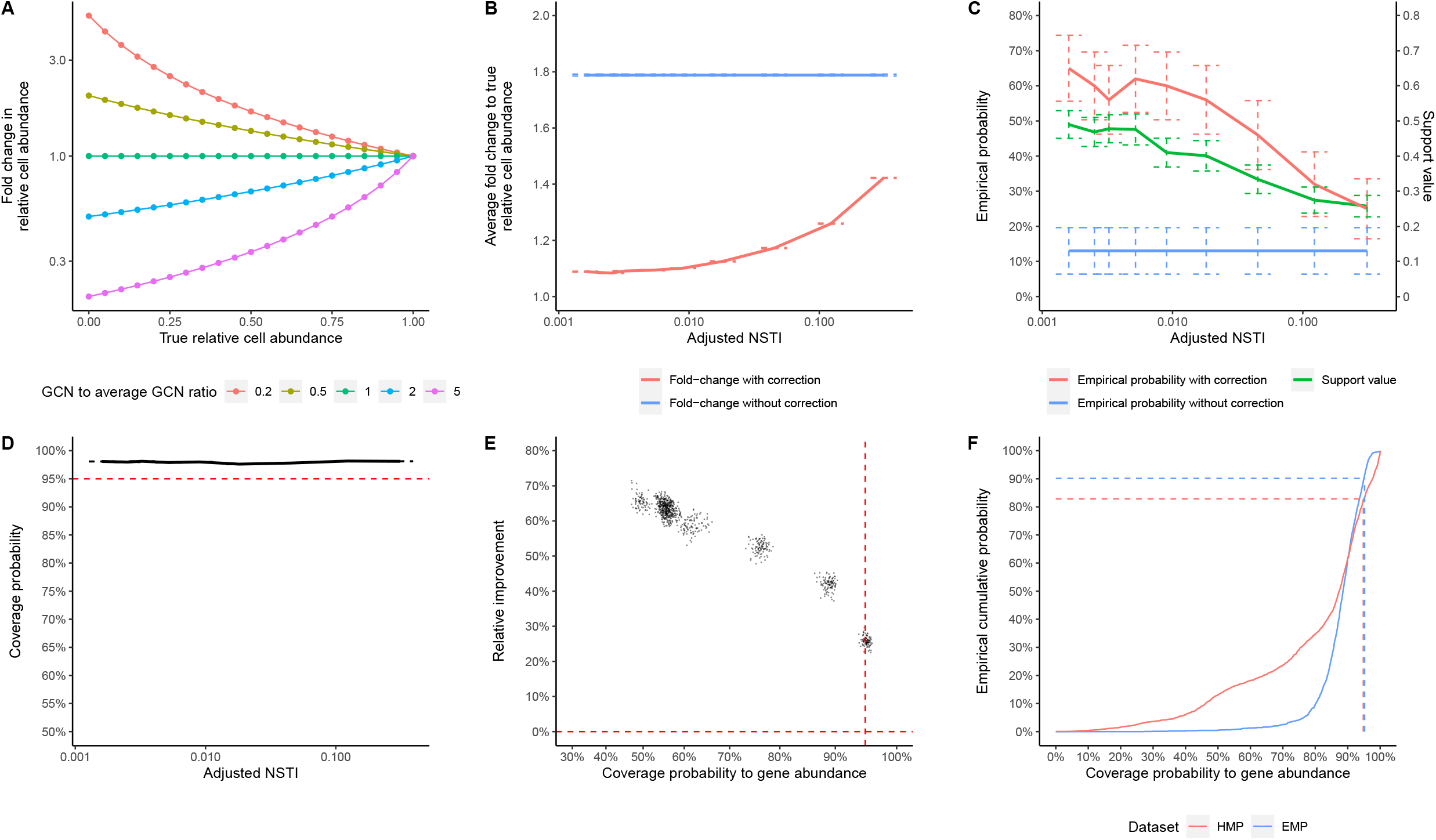
The impact of 16S GCN variation on estimated relative cell abundances. (A) The impact of GCN variation on estimated relative cell abundance based on theoretical calculations. The color of the lines denotes the ratio of an OTU’s of GCN to the average GCN of the community. (B) The average fold-change to the true relative cell abundance. (C) The empirical probability of correctly identifying the most abundant OTU in the community and the support value for the most abundant OTU. (D) The coverage probability of relative cell abundances’ estimated 95% CIs to the true relative cell abundance. Accurate confidence estimates (95% CIs) should produce a coverage probability of 95% regardless of the adjusted NSTI (dashed red line). (E) The correlation between the coverage probability to the relative gene abundance and the improvement by GCN correction. The horizontal red dashed line represents no improvement in relative cell abundance estimates; the vertical red dashed line represents 95% coverage probability to the relative gene abundance. The improvement is quantified by the relative reduction in the difference between the estimated and true cell abundances. (F) The empirical cumulative distribution of the coverage probability to the relative gene abundance in 2560 samples from the HMP1 dataset and 1856 samples from the EMP dataset. All error bars represent 95% CI of the mean.

Because we predict each OTU’s 16S GCN with a confidence estimate, we can provide 95% confidence intervals (95% CIs) for their relative cell abundance as well. Ideally, 95% of the true relative cell abundances should be covered by the 95% CIs. Figure 2D shows that the average coverage probability of the true relative cell abundance is about 98% across NSTD cutoffs, indicating that our 95% CIs are slightly over-conservative. Similarly, we can also calculate the coverage probability of our 95% CI to the relative gene abundance. As expected, when the coverage probability to the relative gene abundance increases, the improvement by GCN correction (quantified by the relative reduction in the difference between the estimated and true cell abundances) decreases (Figure 2E), and that when this coverage probability is below 95%, GCN correction always results in strong improvement in relative cell abundance estimates. In empirical studies when the true abundance is unknown, we can use the coverage probability to the relative gene abundance as a conservative statistic to decide if GCN correction for a community will likely improve the relative abundance estimation or not. For the most abundant OTU in the community, we can calculate its support value from the 16S GCN’s confidence estimates. We found that the calculated support value matches the empirical probability that the most abundant OTU is correctly identified (Figure 2C).

To demonstrate the effect of 16S GCN correction in empirical data, we analyzed the data from the first phase of the Human Microbiome Project (HMP1) and the 2000-sample subset of Earth Microbiome Project (EMP). We found that on average the relative cell abundance with and without 16S GCN correction changes around 1.3-fold in HMP1 and 1.6-fold in EMP. Since the true abundance of OTUs is unknown, we use the coverage probability of 95% CIs to the relative gene abundance described above to evaluate the effect of GCN correction. Our results indicate that a majority of HMP1 (over 82%) and EMP (over 90%) samples have a coverage probability below 95% (as shown in Figure 2F). Since our simulations demonstrate that GCN correction improves the accuracy of relative cell abundance estimation in samples with coverage probability less than 95% (as demonstrated in Figure 2E), this suggests that GCN correction will likely improve relative cell abundance estimates in these HMP1 and EMP samples. In terms of the most abundant OTU, we found that the identity of the most abundant OTU changes after copy number correction in around 20% and 31% of the communities in HMP1 and EMP respectively. The support values for the most abundant OTUs are around 0.85 on average in both datasets, indicating high confidence in the identification of the most abundant OTUs.

### Copy number correction provides limited improvements on beta-diversity analyses

To examine the effect of 16S GCN variation on beta-diversity analyses, we simulated communities at different turnover rates in two types of environments where 0.25%, 1% or 5% of the OTUs are enriched in one environment compared to the other (the SC2 dataset). We found that when the relative gene abundance is used to calculate the Bray-Curtis dissimilarity or the Aitchison distance, the positions of the samples in the PCoA plot shift from their positions based on the true relative cell abundance (solid lines in Figure 3A and B), although this shift is much smaller if the Aitchison distance is used. Correcting for 16S GCN reduces about 56% and 85% of the shifts in the Bray-Curtis dissimilarity (P<0.001, paired t-test, Figure 3A) and Aitchison distance spaces (P<0.001, paired t-test, Figure 3B) respectively. Despite the shift in the PCoA plot, we found that the clustering of communities does not seem to be affected by the 16S GCN variation.

**Figure 3.**
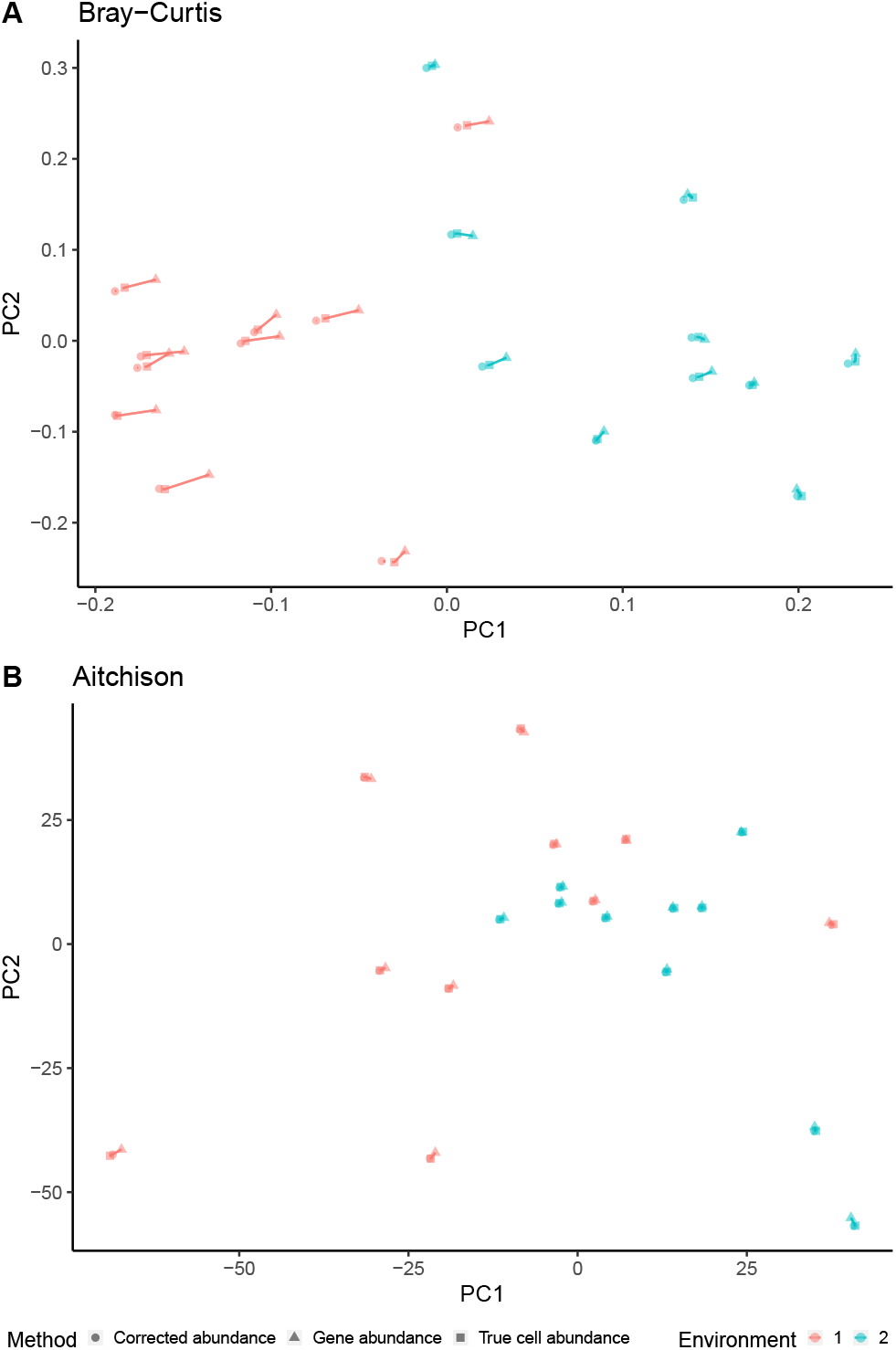
The impact of 16S GCN variation on beta-diversity. Examples of shift in the Bray-Curtis dissimilarity (A) and the Aitchison distance (B) matrices due to 16S GCN variation. The shift for each metric is visualized in a PCoA plot comparing 20 simulated samples from two hypothetical environments with 5 signature OTUs (0.25%) in each environment and a turnover rate of 20%. Solid lines represent the shift of a sample from its true location when using the gene abundance. The results with 1% and 5% enriched signature OTUs are similar to the examples shown in Figure 4.

We observed a limited effect of 16S GCN variation on PERMANOVA. Depending on the metric used, the signature OTU numbers and turnover rates, the proportion of variance explained (PVE) by the environmental type using the true cell abundances ranges from 5.27% to 17.20% on average. Using gene abundance, the average PVE ranges from 5.27% to 17.22% and the change in PVE is not statistically significant regardless the metric used, the signature OTU numbers, or the turnover rates (P>0.002, paired t-test with Bonferroni correction, α=9.26×10^−4^, Table S4), indicating that PERMANOVA is not very sensitive to 16S GCN variation.

It is a common practice to compare the relative cell abundance of OTUs of interest between environments. We found that such comparison is also not sensitive to 16S GCN variation (Table S4), with the fold-change of relative cell abundance estimated using the gene abundance and the truth highly concordant (R^2^ > 0.99). Random forest identified from 20.0% to 89.0% of the signature OTUs when the true cell abundances were used (Table S4). When the gene abundances were used, this recovery rate changes to from 18.0% to 89.32% (Table S4), and the change is not statistically significant (P > 0.032, paired t-test with Bonferroni correction, α=1.85×10^−3^). Correcting for 16S GCN changes the recovery rate to from 17.8% to 89.2% (Table S4), and the change is not significant either (P>0.041, paired t-test with Bonferroni correction, α=1.85×10^−3^). Similar results were found when we examined the effect of 16S GCN variation correction on beta-diversity in empirical data (Supplementary Results).

### Vast majorities of bacterial community studies should benefit from copy number correction

To examine if analysis of real communities would benefit from 16S GCN correction, we calculated the adjusted NSTI for 113842 communities in the microbiome resource platform MGnify (formerly known as EBI Metagenomics) [32] that passed our quality control. These microbiomes were sampled from various environments and include host-associated microbiomes in animals and plants and free-living microbiomes in soil and aquatic environments (Table S5). The adjusted NSTI varies greatly among samples and the median across all samples is 0.01 substitutions/site. In the simulated communities, we observed that GCN correction significantly improves the estimated relative cell abundances (P<0.001, paired t-test) even when the adjusted NSTI reaches 0.3 substitutions/site. We found that more than 99% of the communities from MGnify have an adjusted NSTI less than 0.3 substitutions/site, suggesting that they should benefit from 16S GCN correction when estimating the relative cell abundances. The distribution of adjusted NSTI varies among different environmental types (Figure 4), but the proportion of communities that will likely benefit from 16S GCN correction remains high, ranging from 98% to 100%.

**Figure 4.**
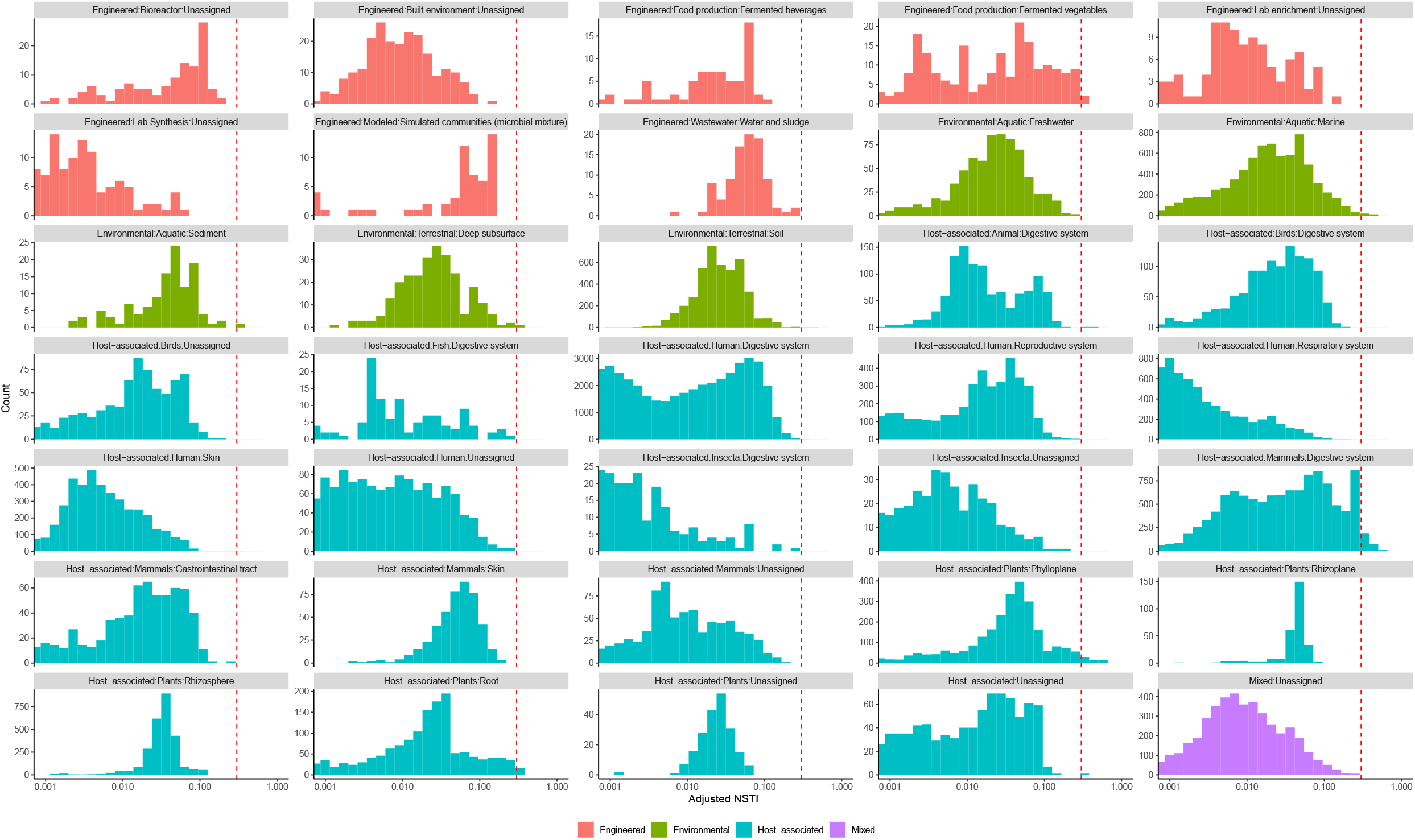
The distribution of adjusted NSTI in empirical data. The distribution of adjusted NSTI of 113842 communities in the MGnify database representing various environmental types. The red dashed line marks the adjusted NSTI of 0.3 substitutions/site.

## Discussion

We address the inherent uncertainty problem in 16S GCN prediction by directly measuring it with confidence estimates. Using simulations and cross-validation, we show that the PE method implemented in *RasperGade16S* outperforms other methods in both the precision and recall rates. This method’s strength comes from three features of its modeling of the 16S GCN evolution: implementation of a pulsed evolution model and accounting for the rate heterogeneity and time-independent trait variation. Pulsed evolution model expects no trait changes to occur over a short branch as jumps are not likely to happen on that branch. This leads to a higher confidence to 16S GCN prediction with a short NSTD, and thus improves the recall of the accurate predictions. By incorporating rate heterogeneity, we can make predictions in the slowly-evolving groups with high confidence, even when their NSTDs are large, thereby further improving the overall precision and recall rates. In the reference phylogeny, 48% of branches were estimated to fall within this slowly-evolving group, whose evolution rate is 145 times slower compared to that of the regularly-evolving group. The third source of improvement for *RasperGade16S* comes from accounting for time-independent variation, which can result from measurement error and intraspecific variation. We show that failing to account for time-independent variation results in model misspecification (Table 1) and overestimated rate of evolution for the pic method.

Having confidence estimates is critical in the presence of inherent uncertainty because they provide direct evaluation of the uncertainty associated with the predictions. Using cross-validation, we show that *RasperGade16S* has high precision (around 0.96), which means for predictions with high confidence (≥95%), 96% of the predictions are accurate. Therefore, we can use the confidence score provided by *RasperGade16S* to select high-quality predictions if necessary, or we can draw firm conclusions from the 16S rRNA data when the confidence is high.

The application of confidence estimation extends beyond the prediction of 16S GCN. Because the uncertainty in the prediction is inherited by statistics derived from the predicted 16S GCN, we can estimate the uncertainty and confidence intervals of important parameters in downstream analyses, such as the relative cell abundance. With confidence intervals, we can draw more meaningful and sound conclusions, such as identifying the most abundant OTU in the community with a support value. Getting confidence estimates of the relative cell abundance is also important for predicting the functional profile of a community based on 16S rRNA sequences. Although PICRUST2 uses an extremely lenient NSTD cut-off to eliminate problematic sequences, it does not provide an accurate confidence measurement of its predictions. As shown in this study, the default maximum parsimony method used by PICRUST2 to predict 16S GCN essentially assumes there is no uncertainty in the predictions, which is unrealistic and leads to poor precision. Incorporation of a more meaningful confidence estimate of 16S GCN prediction in PICRUST2 should make its functional profile prediction more informative.

Strikingly, 99% of 113842 bacterial communities we examined have an adjusted NSTI less than 0.3 substitutions/site, a range where we show that GCN correction improves the accuracy of the relative cell abundance estimation (Figure 2B). Because these communities represent a comprehensive and diverse list of natural and engineered environments, we recommend applying 16S GCN correction to practically any microbial community regardless of the environmental type if accurate estimates of relative cell abundance are critical to the study. Our results therefore affirm the conclusion of the previous studies based on analyses of a much smaller number of communities [5, 7].

Few studies have investigated to what extent the bias introduced by 16S GCN variation will have on the microbiome beta diversity analyses. We show that the effect sizes of 16S rRNA bias on beta-diversity analyses are small. Correcting 16S GCN provides limited improvement on the beta-diversity analyses such as random forest analysis and PERMANOVA test. One possible reason is that for an OTU, the fold change in the relative cell abundance between samples remains more or less the same with or without correcting for the copy number. For example, assuming the estimated relative cell abundances of an OTU in samples A and B are *r_a_* and *r_b_* respectively without copy number correction. When correcting for the copy number, its relative abundance is adjusted with the scaling factor ACN/GCN, where the GCN is the 16S rRNA copy number of the OTU and the ACN is the average copy number of the sample. Assuming the ACN does not vary much between samples, then the scaling factor for the OTU will be roughly the same in samples A and B. So even with copy number correction, the relative abundance change will still be close to *r_a_/r_b_*.

It should be noted that having a confidence associated with the 16S GCN prediction helps to estimate the uncertainty of the prediction, but it does not improve the accuracy of the prediction. Accuracy of the prediction is constrained by the inherent uncertainty, which can only be improved by better sampling the reference genomes. However, as our current sampling is inadequate for accurate 16S GCN prediction of all environmental bacteria, we believe that incorporating confidence estimates is the best practice to control for the uncertainty in the 16S rRNA based bacterial diversity studies, as opposed to not correcting the GCN bias as previously suggested [8, 12].

## Acknowledgements

None.

## Data Availability Statement

The NCBI accession numbers of the reference genomes, the representative 16S rRNA sequences and alignments, the reference phylogeny, the predicted GCN for OTU99 in the SILVA database, the simulated bacterial community data and scripts to reproduce the figures and tables in this study are available in the Dryad repository (https://datadryad.org/stash/share/OaS9BjM_kIVdJ3WkZRT7KO8fDr8D4k8jy3LsOtlYELM). The R package *RasperGade16S* can be downloaded from https://github.com/wu-lab-uva/RasperGade16S. The scripts to conduct the analyses in this study are available in the GitHub repository (https://github.com/wu-lab-uva/16S-rRNA-GCN-Predcition).

