## Supplementary Materials for "Accounting for 16S rRNA copy number prediction uncertainty and its implications in bacterial diversity analyses"

**Supplementary Methods**

16S rRNA GCN quality control

For genomes with multiple copies of 16S rRNA gene, we aligned the 16S rRNA sequences using MAFFT [1] (with parameters: --maxiterate 1000 --globalpair) and picked the 16S rRNA gene sequence that has the highest average similarity (calculated as the proportion of identical bases in the alignment) to other 16S rRNA gene sequences in the genome as the representative sequence. To remove potential errors introduced by mis-assembled genomes [2], we removed genomes whose 16S rRNA GCN differs from their 5S rRNA GCN by greater than 2 copies, genomes whose 16S rRNA sequence contains ambiguous bases, or genomes on the list of withheld genomes in the curated ribosomal RNA operon copy number database rrnDB [3]. The 17 genomes in the rrnDB withheld list are rejected from rrnDB because their 16S rRNA genes are missing, the 16S rRNA GCNs are too high, or the genomes have inconsistent meta data (<https://rrndb.umms.med.umich.edu/withheld/>).

Reconstruction of the 16S rRNA phylogeny

We aligned the remaining representative 16S rRNA gene sequences using HMMER version 3.2 [4] (hmmalign with parameters: --trim --dna –mapali) with the hidden Markov model (HMM) built from the GreenGenes 13.8 16S rRNA gene alignment (hmmbuild with default parameters), and trimmed the alignment with a mask from the GreenGenes database [5]. The HMM, profile alignment and the alignment mask are included in the R package RasperGade16S. After collapsing identical 16S rRNA alignments, 6408 representative sequences remained. They serve as the reference sequences and their taxonomies of are summarized in Table S1. We built a reference tree from the trimmed alignment using RAxML version 8.2 [6] with options -f d -m GTRGAMMA. We used the Deinococcus-Thermus group to root this reference phylogeny. To examine the effect of sequence alignment on model fitting, we also used the 16S rRNA HMM profile from the software Barrnap [7] to align the 16S rRNA genes (hmmalign with default parameters). We trimmed the alignment using a consensus posterior probability threshold of 0.95 (esl-alimask with parameters: -p --ppcons 0.95) and made a 16S rRNA phylogeny as described above.

Modeling 16S rRNA GCN evolution with homogeneous and heterogeneous pulsed evolution models

Using the R package *RasperGade* [8], we fitted one PE model to the entire reference phylogeny and calculated the likelihood of this homogeneous PE model. An analysis of the variance of the PICs associated with each genus indicated that there is a slowly-evolving group and a regularly-evolving group, with the average rate of the slowly-evolving group estimated to be at least 100-fold lower than that of the regularly-evolving group (Figure S1). To model the rate heterogeneity, we created two PE models: *PE_regular_* for the regularly-evolving group and *PE_slow_* for the slowly-evolving group. We then use a two-step iterative binning procedure to estimate the parameters of *PE_regular_* and *PE_slow_* (i.e., jump size and frequency). The *PE_regular_* model was initiated to take the parameter values of the homogeneous PE model. *PE_slow_* was initiated to have a jump size equal to that of *PE_regular_* but a jump frequency100-fold lower. In our first round of binning, from the root to the tip of the reference phylogeny, we classified each node into the regularly- or slowly-evolving group by testing which model (*PE_regular_* or *PE_slow_*) provided a better fit. We merged neighboring nodes belonging to the same group into one neighborhood and flipped neighborhood assignment if the flip resulted in an improved overall AIC value. After the first round of binning, we updated *PE_regular_* and *PE_slow_* by fitting *PE_regular_* to nodes that were classified as regularly-evolving and *PE_slow_* to slowly-evolving nodes. We used the updated models to perform a second round of binning to assign each node in the phylogeny to a group. Finally, we calculated *r*, the rate of evolution in each group, as the process variance per unit branch length defined in a previous study [9]. We then rescaled the reference tree by multiplying the branches in the slowly-evolving group by the ratio *r_slow_*/*r_regular_*. To accommodate time-independent variation in the tip trait values, we calculated a branch length over which the process variance of the fitted pulsed evolution model is equal to the model’s time-independent variation, and added this branch length to each tip branch. We compared the homogeneous and heterogeneous PE models by AIC.

Simulating bacterial communities with 16S rRNA GCN variation

To evaluate the effect of 16S rRNA GCN correction on bacterial diversity analyses, we simulated two sets of bacterial communities using the reference genomes: one set for relative cell abundance analyses (SC1) and the other set for beta-diversity analyses (SC2). We treated each reference genome as one OTU. For SC1, we simulated a total of 100 communities. For each simulated community, we randomly selected 2000 OTUs from the reference genomes, and assigned each OTU a cell abundance randomly drawn from a log-series species abundance distribution.

In SC2, we simulated communities in two environments to evaluate the effect of 16S rRNA GCN correction on beta diversity analyses. We simulated 10 communities per environmental type and 2000 OTUs per community. We controlled the community turnover rate by controlling the number of unique OTUs in each community. For example, at a turnover rate of 10%, a community would have 200 unique OTUs and 1800 core OTUs that are shared among all communities. We varied the turnover rate from 10% to 90% at 10% intervals. To control for the effect size of environmental type, we assigned 5 (0.25%), 20 (1%) or 100 (5%) signature OTUs to each environmental type. These signature OTUs were shared between the two environmental types but were twice more likely to be placed in top ranks of the log-series distribution (i.e., to be more abundant) than the non-signature OTUs in their corresponding environment. The 16S rRNA GCN of each OTU was assigned randomly from the reference genomes’ GCN. We simulated 50 batches of communities for each combination of turnover rate and signature OTU number, resulting in 27000 simulated communities in SC2.

Evaluating the effect of GCN correction in HMP1 and EMP dataset

To check the effect of 16S rRNA GCN correction in empirical data, we analyzed the 16S rRNA V1-V3 amplicon sequence data of the first phase of Human Microbiome Project (HMP1) [10] and the sequence data processed by Deblur [11] in the first release of the Earth Microbiome Project (EMP) [12]. The 16S rRNA GCN for each OTU in the HMP1 and EMP datasets was predicted using *RasperGade16S*. We picked 2560 samples in the HMP1 dataset with complete metadata and used the 2000-sample subset of EMP, and determined the adjusted NSTI and relative cell abundance in each community as described above. For beta-diversity, we randomly picked 100 representative samples from each of the 5 body sites in the HMP1 dataset and analyzed their beta-diversity as described above. For the EMP dataset, we analyzed the beta-diversity within each level-2 EMP ontology (EMPO) category (around 400 to 600 samples per category).

Predicting 16S rRNA GCN for SILVA OTUs

We downloaded 592605 full-length representative bacterial 16S rRNA sequences of non-redundant OTUs at 99% similarity (OTU99) in the SILVA release 132 [13]. We aligned and trimmed the sequences using the method described above. We then inserted the OTUs into the reference phylogeny using the evolutionary placement algorithm (EPA-ng) [14] with the model parameters estimated by RAxML when building the reference phylogeny. We limited the maximum number of placements per SILVA representative sequence to 1. We predicted the 16S rRNA GCN for each SILVA OTU99 as described above using the heterogeneous pulsed evolution model and calculated adjusted NSTDs.

**Supplementary Results**

NSTD values in the context of taxonomy ranks

As NSTD of the 16S rRNA gene also depends on the sequence alignment and how it is trimmed, it can vary from study to study using the same set of reference sequences. Therefore, to put NSTD values of this study in a taxonomic context, we calculated the NSTDs between taxa at different taxonomical levels. For example, at the species level, we calculated the NSTD of a species to another species within the same genus. We found that the median NSTD between congeneric species is around 0.01 substitutions/site and the maximum NSTD threshold (0.464 substitutions/site) in our cross-validation experiment roughly correspond to a taxonomic distance somewhere between class and order (Figure S3).

Copy number correction provides limited improvements on beta-diversity analyses in empirical data

We analyzed the beta-diversity using the HMP1 and EMP datasets. Because we observed that the effect of GCN correction is independent of the metric used in beta-diversity analyses, we only used Bray-Curtis dissimilarity in HMP1 and EMP datasets. We found that correction of 16S rRNA GCN does not seem to affect the clustering of communities by body sites in the HMP1 PCoA plot. Pairwise PERMANOVA shows that the mean PVE by the body site in HMP1 is 14.9% before 16S rRNA GCN correction and decreases marginally to 14.6% after correction, and the PVEs using the gene abundance and the corrected cell abundance are also highly concordant (R^2^>0.98). In EMP, within each level-2 environment (EMPO2), the average PVE by level-3 environment (EMPO3) remains at 7.7% before and after 16S GCN correction and the PVEs using the gene abundance and the corrected cell abundance are highly concordant (R^2^>0.99) as well. On the other hand, pairwise random forest tests yield similar results before and after 16S rRNA GCN correction, with around 9 out of the top 10 features identified by the random forest test remaining unchanged before and after correction in HMP1 and around 8 out of the top 10 unchanged in EMP. In terms of the fold-change of relative cell abundances between body sites, we found that copy number correction has little impact as the estimated fold-change before and after correction are highly similar (R^2^>0.95) in both datasets.

Predicting 16S rRNA GCNs for SILVA OTUs

Using *RasperGade16S*, we predicted the 16S rRNA GCN for 592605 bacterial OTUs (99% identity) in the release 132 of the SILVA database. Overall, the median adjusted NSTD for all bacterial OTUs is 0.070 substitutions/site, and 34.7% of the predictions have a high confidence of 95% or greater, and 74.9% of the predictions have a moderate confidence of 50% or greater (Table S6). This shows that for most OTUs in the SILVA database, the phylogenetic distance to a reference 16S rRNA is small enough that we can have reasonable confidence in the predictions. In comparison, randomly guessing has a null confidence of around 6.7% (1 out of 15 possible GCNs). Among major phyla with more than 10000 OTUs, the proportion of highly confident predictions varies greatly (Table S6), with Cyanobacteria having the lowest proportion of 19.1% and Acidobacteria having the highest proportion of 50.4%. Similarly, the proportion of moderately confident predictions varies from 58.3% to 89.5% among these phyla. Interestingly, the proportions of highly confident predictions closely match the proportions of slowly-evolving OTUs in each phylum (Table S6), suggesting a causal relationship between them.

**Figure S1. The distribution of local rate of 16S rRNA GCN evolution.** The distribution of local rate of 16S rRNA GCN distribution among bacterial genera, estimated from the variance of the PICs. A small rate of 0.1 is added to the estimated rates for displaying the rate in the logarithmic scale. The red dashed line divides the rates into two groups: slow-evolving group (left) and regularly-evolving group (right).

**Figure S2. The distribution of rate groups along the 16S rRNA reference phylogeny.** The distribution of rate groups is denoted by colors. Red color represents slowly-evolving group and black color represents regularly-evolving group. The branch lengths displayed in the figure are not scaled by the GCN evolution rate.

**Figure S3. The distribution of NSTD at different taxonomic levels.** The solid horizontal lines inside the violin plot represent the quartiles of the distribution. The red dashed line represents the maximum NSTD threshold used in the cross-validation experiments. The unit of NSTD is the same as the unit of branch length of the reference phylogeny (substitutions per site).

**Table S1. The taxonomic composition of genomes in the reference phylogeny.** The count of reference genomes within a clade is listed at phylum, class, order, and family level.

**Table S2. The effect of HMM profiles on model fitting**. The AIC and parameters of fitted Brownian motion (BM) and pulsed evolution (PE) models when the alignment of 16S rRNA genes uses different HMM profiles are listed.

**Table S3.** **Fitted parameter of homogeneous and heterogeneous pulsed evolution models**. The jump frequency, jump size and the magnitude of time-independent variation of the fitted homogeneous and heterogeneous pulsed evolution models are listed. The unit of jump frequency is jump per unit branch length.

**Table S4.** **The effect of 16S rRNA GCN correction on beta-diversity analyses.** The performance statistics of random forest tests, PERMANOVA, and abundance comparison before and after 16S rRNA GCN correction are listed at different signature OTU number and turnover rate.

**Table S5.** **Count summary of environmental types in MGnify dataset.** The count number of samples within each biome (environmental type) is listed at the first and second biome level.

**Table S6.** **Summary of SILVA 16S rRNA GCN predictions.** Highly confident predictions are defined as predictions with a confidence of 95% or greater. Moderately confident predictions are defined as predictions with a confidence of 50% or greater.
