## Supplementary figures and images for "Accounting for 16S rRNA copy number prediction uncertainty and its implications in bacterial diversity analyses"

### Fig S1

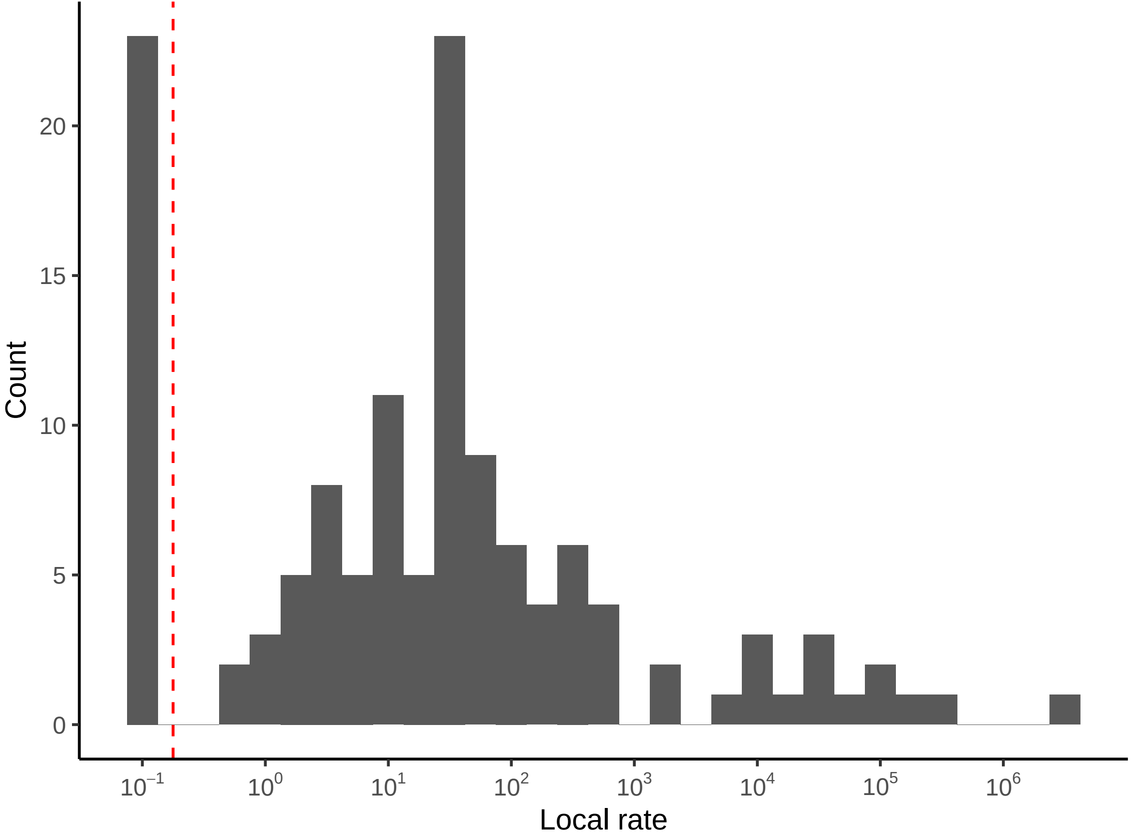

### Fig S2

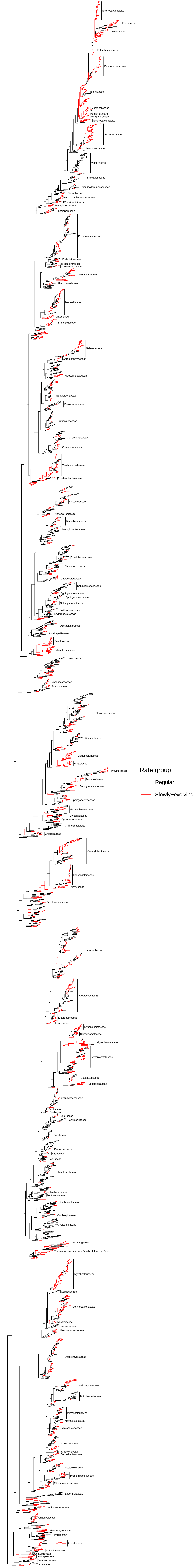

### Fig S3

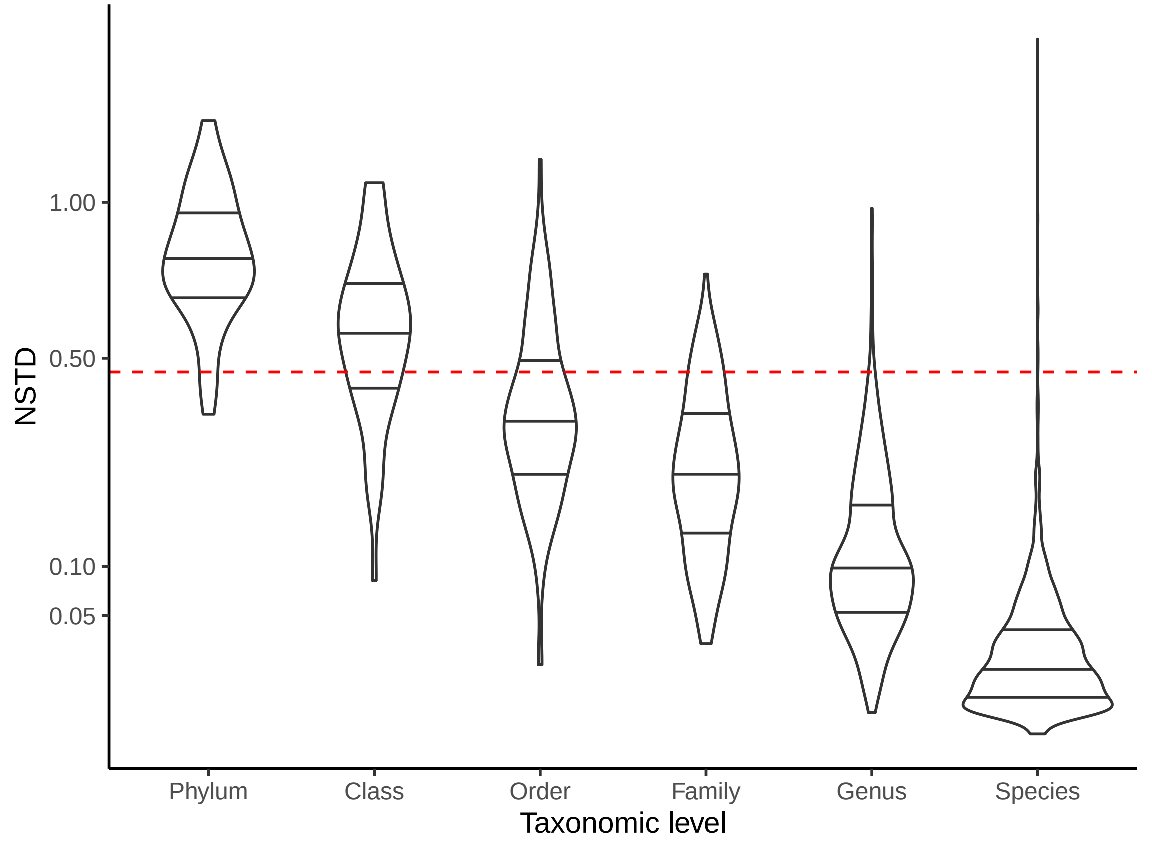
